# Genular: An Integrated Platform for Defining Cellular Identity and Function through Single-Cell Gene Expression and Multi-Domain Biological Data

**DOI:** 10.1101/2024.10.23.619445

**Authors:** Ivan Tomic, Stephanie P Hao, Adriana Tomic

**Affiliations:** Department of Virology, Immunology & Microbiology (VIM), Boston University; Department of Biomedical Engineering (BME) & Virology, Immunology & Microbiology (VIM), Boston University

**Keywords:** single-cell RNA-seq integration, cell marker score calculation, immune cell heterogeneity, gene signature identification, cellular differentiation, systems immunology, gene expression analysis, immunogenomics

## Abstract

Accurately defining cellular identity and function is essential for advancing immunology, understanding disease mechanisms, and developing targeted therapies. However, current bioinformatics tools are limited in their ability to integrate and analyze the vast and diverse single-cell RNA sequencing (scRNA-seq) datasets available, hindering the comprehensive capture of cellular heterogeneity and the identification of subtle genetic changes across immune states, differentiation pathways, and tissue contexts. To overcome these challenges, we introduce *genular*, an open-source platform that unifies gene expression data analysis across diverse cell types by integrating scRNA-seq data with extensive genomic and proteomic information from 16 databases, including NCBI Gene, Human Protein Atlas, STRING, and UniProt. *genular* aggregates data from more than 2,893 scRNA-seq experiments, encompassing over 74.5 million unique cells across various tissues and conditions. A key feature of *genular* is calculating a cell marker score for each gene, enabling the quantification of gene expression across cells to derive unique profiles specific to cell types, states, and lineages. Using *genular*, we differentiate T cell memory states, map differentiation profiles by tracking gene expression changes as monocytes mature into macrophages and lymphoid progenitor cells develop into T cells, and capture tissue-specific reprogramming of macrophages, revealing distinct gene expression profiles that enable specialized functions in different tissues. By integrating scRNA-seq data with multi-domain biological information and employing advanced statistical methodologies, *genular* provides a scalable platform that accurately defines cellular identities, functional states, and differentiation pathways. This comprehensive approach facilitates breakthroughs in immunology, gene regulation, cellular differentiation, and disease research, enabling a deeper understanding of immune cell functions and their roles in health and disease.

## Introduction

In immunology, the cell is the fundamental unit through which immune responses are orchestrated, arising from complex interactions among various cell types that each perform specific functions to defend the body against pathogens. Traditionally, immune cells such as T cells, B cells, macrophages, and dendritic cells have been classified based on size, morphology, and the presence of proteins, i.e., cellular markers identified through microscopy and flow cytometry [47]. However, these cellular markers represent only a small fraction of the cellular genome, failing to capture the full heterogeneity of the immune system. Immune cells within the same lineage can undergo subtle genetic changes that drive distinct states and functionalities across different tissues and phases of the immune response, affecting how they respond in various physiological contexts [26]. For example, memory T cells are divided into effector-memory (EM) and central-memory (CM) subsets, each playing unique roles in immune protection: EM T cells provide rapid protection in peripheral tissues by executing immediate effector functions, whereas CM T cells reside in lymphoid organs and generate proliferative responses to combat systemic pathogens [16, 53, 34]. Accurately defining cell identity based on genetic programming is crucial for understanding these diverse immune responses and their regulation. However, several critical research questions and gaps remain in accurately characterizing and tracking cellular identities to enhance our understanding of disease pathogenesis and develop targeted therapies. While there has been progress in areas like cancer immunotherapy by identifying exhausted T cells versus functional effector T cells to inform treatment strategies [68], comprehensive methods for defining and utilizing cellular biomarkers across diverse immune environments are still lacking. Additionally, understanding the dynamic changes in gene expression as immune cells migrate between tissues and adapt to different physiological contexts poses a significant challenge [47]. These gaps highlight the need for more robust and scalable approaches to accurately define cellular identities and their functional states within the immune system, ultimately improving our ability to diagnose, treat, and prevent immunological diseases.

Advances in microfluidics and improvements in RNA isolation and amplification methods have enabled the isolation and profiling of large numbers of immune cells at the single-cell level using next-generation sequencing technologies. High-throughput single-cell RNA sequencing (scRNA-seq) offers unprecedented insights into cellular heterogeneity by capturing gene expression profiles of individual cells [62]. Large-scale initiatives, such as the Human Cell Atlas [50], the NIH Human BioMolecular Atlas Program (HuBMAP) [21], and the LifeTime [49], aim to characterize human tissues and cell types comprehensively. However, these repositories primarily focus on cataloging cell types and states without integrating this information with other biological databases, limiting their capacity to leverage cell information as functional units within biological pathways and networks.

Moreover, common computational strategies for analyzing scRNA-seq data involve unsupervised machine learning techniques [9], such as t-distributed stochastic neighbor embedding (t-SNE) [66] and uniform manifold approximation and projection (UMAP) [4], to manage high dimensionality and reduce noise. While these techniques enable visualization and clustering of cells into distinct populations, they are computationally intensive and infeasible for the vast number of cells available in public repositories, exceeding 74.5 million. Additionally, clustering can obscure subtle differences and assign gene signatures at the subpopulation level rather than full single-cell resolution [25]. Consequently, there is a pressing need for statistical approaches that can efficiently analyze large-scale scRNA-seq data without relying on computationally intensive methods.

To address these challenges, we developed *genular*, an open-source platform that unifies and streamlines the analysis of gene expression data across diverse immune cell types by integrating scRNA-seq data with extensive genomic and proteomic information. *genular* leverages the Cell Ontology database to map each cell to standardized cell type definitions, ensuring consistent and standardized annotations facilitating cross-study comparisons. Furthermore, it incorporates gene ontology and functional annotations from resources such as NCBI Gene [8], Human Protein Atlas [65], STRING [60], UniProt [11], and Gene Ontology to provide comprehensive insights into gene functions, pathways, functional annotations, and interactions. Utilizing a novel statistical framework, *genular* calculates cell marker scores for each gene, enabling the quantification of gene expression across all 74.5 million unique cells to derive unique expression profiles specific to cell types, states, and lineages. This approach enhances the detection of subtle cellular states, mitigates technical noise, and supports scalable, high-performance data processing. By allowing researchers to use cell information as functional components within biological pathways, *genular* enables comprehensive exploration of cellular function within complex biological systems, empowering new discoveries in immunology, gene regulation, cellular differentiation, and disease research.

## Materials and methods

### Data Acquisition and Processing

Data acquisition is automated through a series of scripts located in the `data/` directory. The primary script, `download.sh`, facilitates the retrieval of various biological datasets, including gene annotations, protein sequences, and gene expression data. These datasets are sourced from well-established resources such as NCBI Gene [40], Pfam [57], PROSITE [55], Human Protein Atlas [64], Reactome Pathway Knowledgebase [37], STRING [61], and UniProt [12].

Due to the substantial size of these datasets, which collectively exceed 4.8 TB, we have developed a highly efficient, multicore processing pipeline. This pipeline leverages custom Node.js-based file readers, specifically designed to handle large-scale XML (Extensible Markup Language) and JSON (JavaScript Object Notation) files without overwhelming system memory. By reading files sequentially, these readers enable processing of even the largest datasets, while parallelizing the workload across multiple CPU cores.

The acquisition of single-cell expression data, particularly from CELLxGENE [13], is managed separately, requiring custom integration into the pipeline.

### Single-cell data retrieval and pre-processing

Single-cell expression data from CELLxGENE[13, 14] is retrieved using a custom Puppeteer[48] script designed to automate interactions with the CELLxGENE web interface. The retrieval process involves several key steps:

#### Data Retrieval

i. **Automated navigation**: The Puppeteer library is used to control a headless browser, navigating to the CELLxGENE collections page (https://cellxgene.cziscience.com/collectio and scrolling through available datasets to capture their links. NodeJS[17] script is located under ‘/data/unprocessed/cellxgene/index.js’. Since bulk download or programmatic access to single-cell expression files on CELLxGENE[13, 14] is not easily supported, this automated solution efficiently gathers the required data by mimicking manual browsing actions.
ii. **Collection download**: For each collection, the script interacts with the dataset download buttons and opens the modal window to retrieve available files in H5AD[70] format.
iii. **File verification**: Before downloading, the script checks if the dataset already exists locally to avoid redundant downloads.

#### Conversion to JSON

After the single-cell expression data is downloaded from CELLxGENE[1 14] in H5AD[70] format, it is converted into JSON[7] files for integration into the database. The conversion process involves the following steps:

i. **Data loading**: Each H5AD[70] file is loaded into memory using the Scanpy Python library, which facilitates efficient manipulation of single-cell data. During this step, the cell and gene annotations are mapped to their respective ontology IDs.
ii. **Gene expression processing**: For each gene in the dataset, expression levels across all cells are extracted. Non-zero expression values are identified, and the corresponding cell types are recorded. This ensures that gene expression is linked to the specific cell types where it occurs.
iii. **Parallel processing**: To enhance performance, the dataset is split into chunks, and gene expression data is processed in parallel using multiple (95) CPU cores. This parallelization significantly reduces the time required to convert large datasets.
iv. **Output generation**: The processed data is saved in a structured JSON[7] format, with each file named according to the gene, tissue, and disease ontology term IDs. The resulting JSON[7] files capture the relationships between genes and the cell types in which they are expressed, allowing for easy querying and integration into the database.

This conversion process standardizes high-resolution single-cell expression data, making it accessible for further downstream analysis and seamless integration into the database. The Python script responsible for this conversion is implemented in `process_data/main.py` within the `cellxgene` dataset.

#### Processing

Once the JSON files are generated, they are processed to calculate marker scores and other relevant metrics. The ‘combine.uberon.js’ script is used to aggregate data from multiple JSON files and standardize it for further analysis.

The process involves the following steps:

i. **File loading**: The script reads all JSON files from the ‘bulk/’ directory, matching UUIDs[29] using regular expressions to organize them by tissue and disease type.
ii. **Gene data aggregation**: For each gene, the script creates an object structure that links each gene to its tissue, disease, and experiment. This object (‘allGenes’) is progressively populated by iterating through the gene expression values found in the JSON files.
iii. **Cell lineage collection**: The script collects lineage data for each cell by querying external resources like Ontology Lookup Service (OLS)[54] Solr[2] database for parent and child entries, creating a hierarchical structure of cells.
iv. **Expression data processing and marker score calculation**: Each cell’s expression values are standardized, and effect sizes are calculated for marker genes across different cellular contexts (tissues, diseases, experiments and overall). The effect size is defined as the difference between the expression of the gene in the target cell and the median expression in other cells. These effect sizes are then averaged to obtain the marker score for each cell type, providing a quantitative measure of gene overexpression in that cell compared to others.
v. **Data output**: Once all the calculations are complete, the results are written to new JSON files that can be imported to our database. The script saves cell-linked entities, cell lineages, and processed gene data into the ‘output/’ directory.

#### Marker Score Calculation and Threshold Determination

We aim to identify significant gene expression deviations across different cellular contexts by calculating a marker score for each cell type. This score quantifies how much a gene’s expression in a specific cell deviates from the average expression across other cells, highlighting potential biomarkers and key biological functions.

##### Effect Size Calculation

For each gene *g* and each cell type *i*, we collect the expression data *X*_*gi*_. The effect size *E*_*gi*_ measures the difference in gene expression between cell *i* and all other cells *j* not in the same lineage or context. Specifically, we compute the effect size as:

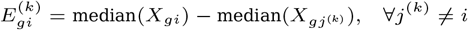

where:

- *X*_*gi*_: Expression data for gene *g* in cell *i*.
- 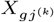 : Expression data for gene *g* in comparison cell *j*^(*k*)^.

This calculation yields a set of effect sizes 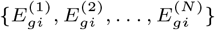 for each cell *i*, where *N* is the number of comparisons made.

##### Marker Score Calculation

To robustly summarize the effect sizes and mitigate the influence of outliers, we employ the *bootstrap percentile method* to calculate the marker score *S*_*gi*_ for each cell *i* within a specific context *c*. The marker score is defined as the 10th percentile of the bootstrapped distribution of effect sizes:

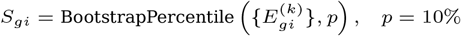

where:

- *BootstrapPercentile*: A method that estimates the percentile *p* of the effect size distribution using bootstrapping with *B* resamples.
- *p*: The desired percentile (in this case, 10%).

###### Bootstrapping Procedure

i. *Resampling*: From the set of effect sizes 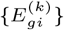, we generate *B* bootstrap samples by randomly sampling with replacement.
ii. *Percentile Calculation*: For each bootstrap sample, we calculate the desired percentile *p*.
iii. *Marker Score*: The average of these percentiles across all bootstrap samples yields the marker score *S*_*gi*_.

This approach provides a robust estimate of the lower end of the effect size distribution, capturing consistent overexpression signals while reducing sensitivity to extreme values.

##### Threshold Determination Using Standard Deviation

To determine the significance o f t he m arker s cores, w e e stablish a threshold *T*_*c*_ for each context *c*. This threshold is calculated using the mean and standard deviation of the marker scores across all cells within the context:

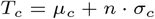

where:

- *T*_*c*_: Threshold for the given context *c* (e.g., tissue).
- *µ*_*c*_: Mean of the marker scores *S*_*gi*_ within context *c*.
- *σ*_*c*_: Standard deviation of the marker scores within context *c*.
- *n*: Number of standard deviations added to the mean (set to *n* = 1 for this analysis).

By incorporating the standard deviation, the threshold accounts for the variability of marker scores, allowing us to identify cells with exceptionally high marker scores.

##### Identification of Significant Cells

Cells with marker scores exceeding the threshold *T*_*c*_ are considered to have significant overexpression of gene *g* in context *c*:

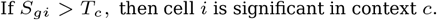

This criterion ensures that only cells with marker scores notably higher than the average are identified, e nhancing t he s pecificity of our analysis.

##### Implementation Details

###### In our code implementation

- **Effect Size Computation**: We calculate the effect sizes by comparing the median expression of each cell to that of other cells, ensuring robustness against outliers.
- **Bootstrapping**: We perform *B* = 250 bootstrap resamples to estimate the 10th percentile of the effect sizes for each cell, providing a reliable marker score.
- **Threshold Calculation**: By using the mean and standard deviation of the marker scores within each context, we set a dynamic threshold that adapts to the distribution of scores, improving detection of significant overexpression.

##### Statistical Justification

- **Bootstrapping Techniques**: Bootstrapping is a non-parametric approach that does not assume a specific distribution of the data, making it suitable for complex biological datasets where normality cannot be guaranteed [15].
- **Standard Deviation in Thresholding**: Incorporating the standard deviation allows the threshold to reflect the dispersion of marker scores, ensuring that only cells with statistically significant deviations are selected [38].

##### Conclusion

By combining bootstrapped percentile marker scores with thresholds based on standard deviation, we provide a robust framework for identifying significant gene overexpression across different cellular contexts. This method enhances the reliability of marker detection, offering valuable insights into gene functions and potential therapeutic targets.

### Data parsing and database population

The ‘parsers/’ directory contains scripts responsible for parsing the processed data and importing it into the MongoDB[23] database. The main script, ‘parsers/main.js’, orchestrates the parsing of all collected database files and populates the respective MongoDB collections.

Key datasets are processed as follows:

- **Gene annotations**: Files such as `gene2accession`, `gene2pubmed`, `gene_info`, `gene2refseq`, and `gene2ensembl` are downloaded from NCBI Gene.
- **Protein sequences**: UniProt protein data is retrieved via the `uniprot_sprot.xml`, `uniprot_trembl.xml`, and `idmapping_selected.tab` files.
- **Gene-protein asociations**: Data from resources like Reactome and Pfam, including files like `ensembl2reactome`, are integrated to map gene-protein interactions.
- **Gene ontology and functional annotations**: Files like `gene2go` are processed to provide gene ontology information.
- **Cellular expression data**: Processed single-cell data from `cellxgene` provides gene expression information in various cell types.
- **Taxonomy data**: The `taxdump` file contains taxonomy information for genes.
- **Gene droup onformation**: Data about gene groups is sourced from the `gene_group` file.
- **Tissue expression data**: Protein Atlas data is retrieved from the `proteinatlas.xml` file to map protein expression across tissues.

Given the substantial size of some datasets, exceeding 500GB, we developed custom “line-by-line” parsers for efficient processing of large XML and JSON files. These parsers, implemented in `linebyline.js`, enable scalable data handling and are an integral part of our data processing pipeline.

### Database Schema

The *genular* database is built using MongoDB[23], a NoSQL database chosen for its flexibility and ability to handle large datasets. At the core of the database is the *gene* object, which is represented by a JSON schema. This schema is flexible, as is typical for NoSQL databases, allowing for scalable and dynamic data storage that accommodates the evolving nature of biological data.

The full schema, detailing all fields and attributes within the gene object, can be accessed on the project website: https://genular.atomic-lab.org. For a visual overview of how the various databases are integrated and processed.

The full JSON schema for the gene data used in this study is available for download. You can download the complete database dump or respective gene schema at *genular* website: https://genular.atomic-lab.org

### Access Methods

*genular* offers three primary access methods: a RESTful API, an R package[27], and a complete MongoDB database dump there is also an online interface https://genular.atomic-lab.org that can be accessed remotely or locally deployed. The MongoDB dump can be imported into a local environment for advanced queries and custom analyses. The R package can be configured with a custom endpoint URL for flexibility in local or remote data access.

#### RESTful API

The RESTful API provides programmatic access to *genular*, enabling seamless integration with computational workflows. It supports a wide range of queries related to genomic data, cellular expression data, and pathway suggestions, making it a valuable tool for researchers and bioinformaticians. The API is structured to offer flexibility in data retrieval, allowing users to access the information they need for their specific research use cases.

The API exposes several endpoints for retrieving gene, cell, and pathway information. It supports both GET and POST requests to ensure versatility in data interaction, allowing both simple retrieval and more complex data filtering and analysis.

The API is built using the Slim framework[31] and offers comprehensive documentation available at https://genular.atomic-lab.org/api-docs/, detailing each endpoint, required parameters, and example responses. Below is a summary of the key features and endpoints:

- **Gene Information:** The API allows users to retrieve detailed gene information, including gene annotations, protein sequences, and associated diseases. The primary route for this is the `/api/v1/gene/search` endpoint, which supports querying based on various gene attributes.
- **Cell Information:** Researchers can retrieve cell-specific data using the `/api/v1/cells/search` endpoint, which provides information on cellular gene expression. Additionally, the `/api/v1/cells/suggest` endpoint offers functionality for cell suggestion based on input queries, allowing researchers to identify relevant cell types based on gene expression patterns.
- **Pathway Information:** Pathway-related data can be accessed via the `/api/v1/pathways/suggest` endpoint, where researchers can retrieve pathway suggestions relevant to specific gene sets or biological processes.
- **Authentication and API Keys:** API access is secured using API keys. This ensures secure and authorized access to the database.
- **OpenAPI Documentation:** The API documentation follows the OpenAPI specification, allowing users to explore available routes and query parameters interactively.

#### R Package

*genular* offers an R package[63], available on GitHub and published on CRAN, that streamlines data access and analysis, designed to simplify querying the database and integrating *genular*’s datasets directly into R-based pipelines. The package provides functions to interact with the API endpoints, handle authentication, and process the returned data structures. Documentation is available on GitHub[27] and additional usage examples are available in the package vignette hosted on CRAN https://cran.r-project.org/web/packages/genular/readme/README.html.

Example 1: Determining Cell IDs and Marker Score Thresholds

The *genular* R package makes it simple to retrieve pre-calculated marker scores and cell-specific thresholds. These marker scores, which quantify how much a gene’s expression deviates from the average, help researchers identify significant gene expression patterns within specific cell types. By leveraging the pre-calculated thresholds stored in the *genular* database, users can quickly pinpoint which genes are significantly overexpressed in particular cells (Supplementary Methods 7.0.1).

Example 2: Retrieving Single-Cell Gene Expression Profiles Across Specific Cell Types

The package also provides a robust querying system for retrieving detailed gene information from the *genular* database. Users can search for genes based on IDs or other criteria, accessing data such as gene symbols, cross-references, and cell-specific expression profiles. This feature is especially useful for researchers investigating gene functions, relationships to diseases, or roles in specific cell types. (Supplementary Methods 7.0.1).

Example 3: Searching for Gene Information

Researchers can retrieve or filter specific gene information, including gene IDs, symbols, cross-references, expression profiles for various cell types, or any other gene-related fields defined in the gene MongoDB schema, available at https://genular.atomic-lab.org. The query searches for genes based on specific IDs, enabling detailed analysis of gene functions and their roles in single-cell gene expression data (Supplementary Methods 7.0.1).

More examples of how to use the *genular* database can be found in package vignette.

#### Database Dump

To support large-scale analyses and offline processing, a full MongoDB database dump is available for download. This allows users to host and query the entire *genular* database locally. The dump can be accessed on *genular* website https://genular.atomic-lab.org.

## Results

### genular architecture

*genular* is a scalable, open-source platform designed for large-scale integration and analysis of single-cell transcriptome data by unifying it with extensive genomic and proteomic information (Fig. 1). At the core of *genular*’s architecture is a robust database schema built with MongoDB, which organizes gene-related data into structured objects encompassing gene identifiers, taxonomy information, gene annotations, and protein interactions. Central to *genular* is the integration of the Cell Ontology database, which maps each cell to standardized ontology term IDs, ensuring consistent cell type definitions and enabling cross-study comparisons (Fig. 1a). *genular* processes high-dimensional gene expression data from scRNA-seq efficiently to derive unique expression profiles across diverse immune cell types and tissue contexts. This is achieved by integrating data from multiple biological databases, such as NCBI Gene[8], Human Protein Atlas[65], STRING[60], and UniProt[11], enabling comprehensive annotation of gene functions, pathways, and protein interactions (Figs. 1b, 1c).

**Fig 1.**
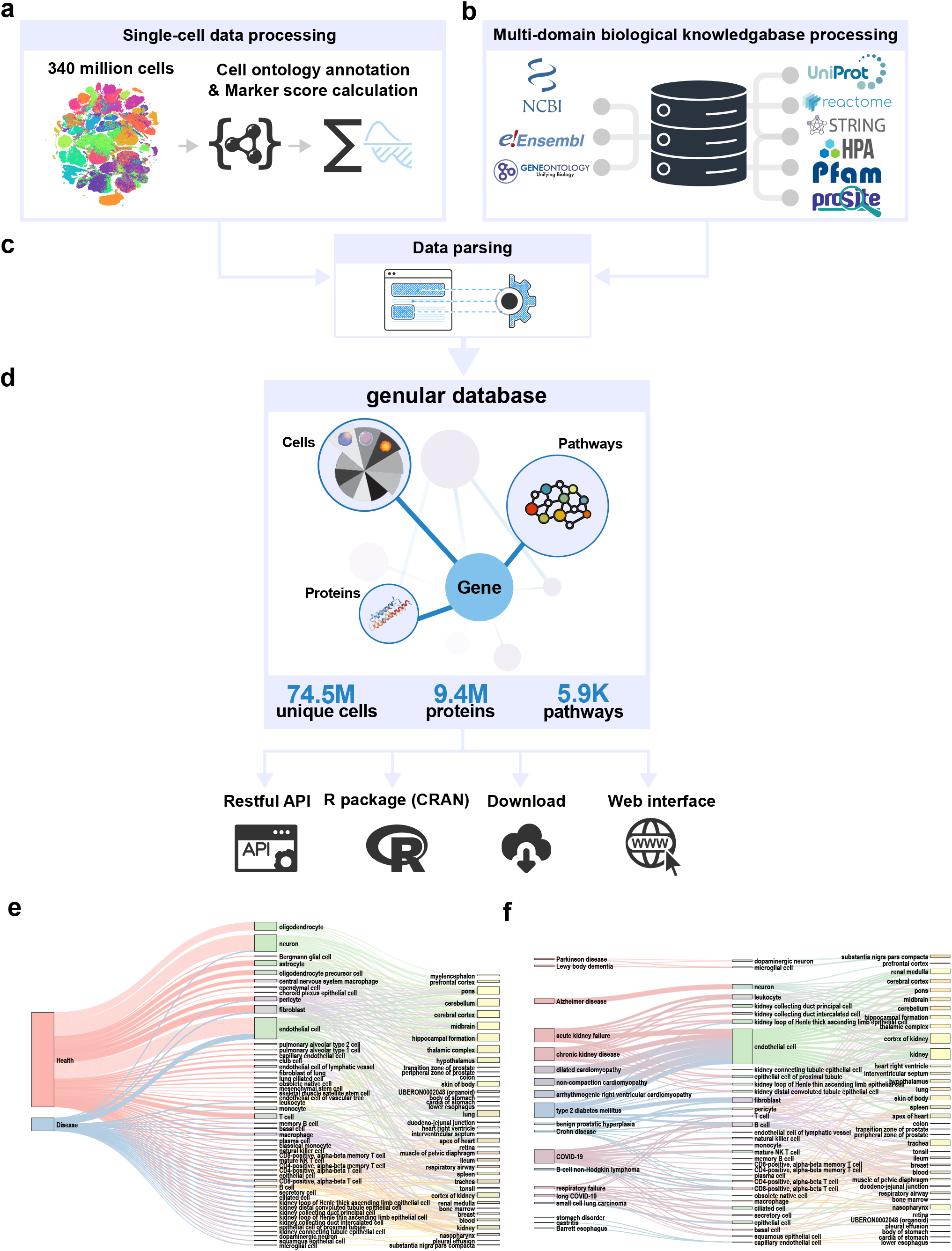
Overview of the *genular* platform’s architecture. (a) Integration of the Cell Ontology for consistent cell type definitions. (b) Biological database integration for gene annotation. (c) Processing single-cell data and multi domain biological databases using data parsing and populating database. (d) genular database contains 74M unique cells connected across 9.4M proteins and 5.9k pathways. It can be accessed through API, R package, data download or web interface. (e,f) Visualization of number of top unique cells across diseases and tissues in genular database subset.

We assembled a large-scale dataset comprising over 74 million unique single-cell transcriptomes across more than 2,893 scRNA-seq experiments (Fig. 1d). Rigorous quality control measures were employed to exclude low-quality or unreliable cells, such as dead or dying cells, doublets, cells with low RNA content, or those affected by technical artifacts, thereby ensuring the integrity of the dataset. *genular*’s architecture is designed for scalability and offering accessibility through a RESTful API, an R package, and local deployment options (Fig. 1d). By integrating scRNA-seq data with multi-domain biological information and employing advanced statistical methodologies, *genular* provides a unified platform for precisely defining cellular identities, functional states, and differentiation pathways across diverse immunological contexts (Figs. 1e, 1f).

### genular Statistical Framework for Gene Expression Visualization from single-cell transcriptome data

*genular* normalizes and processes the gene expression profile of each single cell through a statistical framework that calculates cell marker scores for each gene (Fig. 2). Rather than using rank-based encoding, *genular* directly quantifies deviations in gene expression by comparing each gene’s expression level within a cell type to the average expression level across all cell types within the dataset (Fig. 2a). This method allows for precise identification of upregulated or downregulated genes specific to particular cell types, facilitating the detection of crucial markers without relying on clustering techniques like t-SNE[66] or UMAP[4].

**Fig 2.**
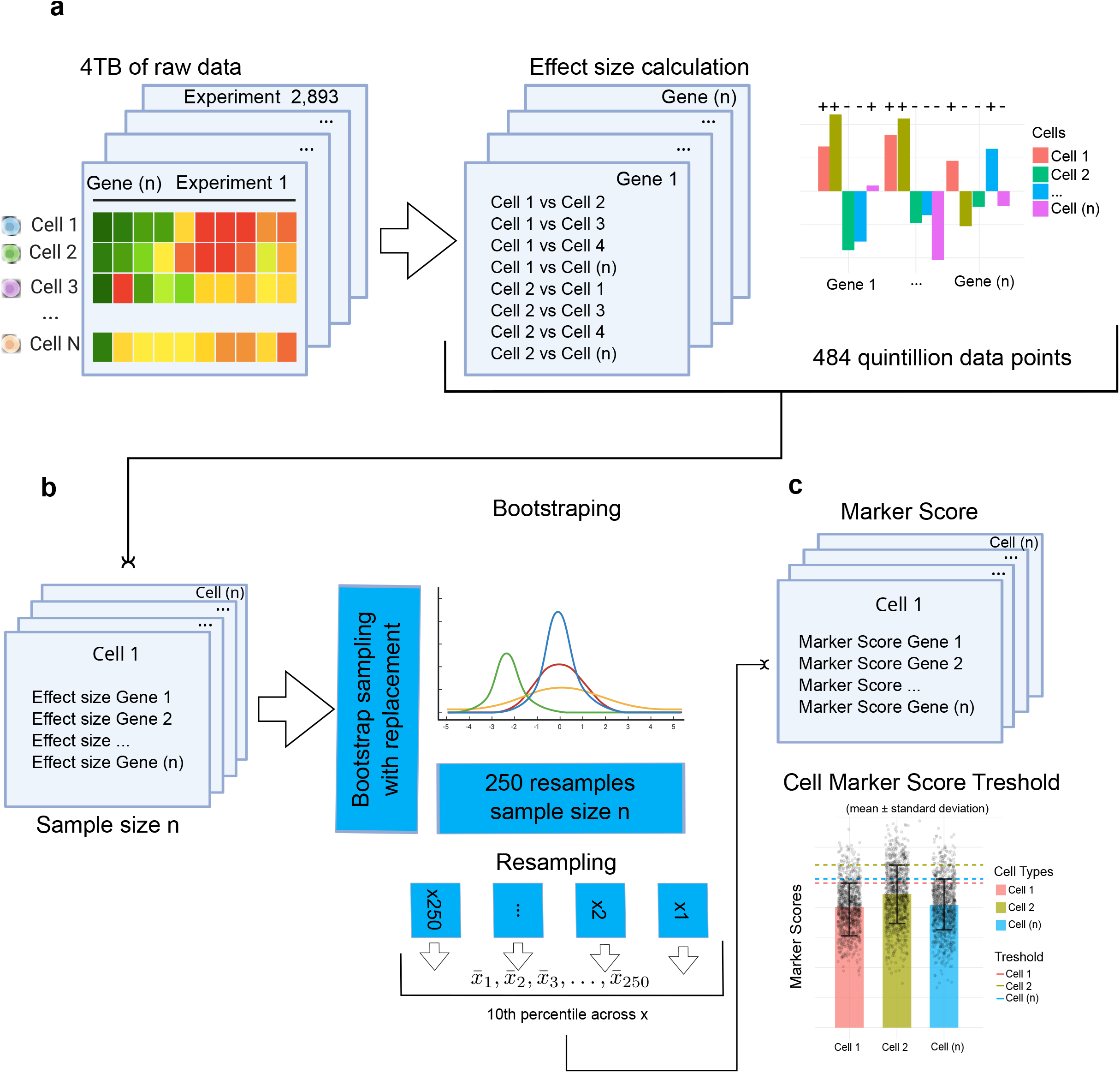
Framework for calculating cell marker scores. (a) Direct quantification of gene expression deviations and effect size calculation, resulting in 484 quintillion data points across cell comparisons. (b) Bootstrapping approach with 250 resamples and 10th percentile calculations to ensure robust marker score estimates. (c) Dynamic determination of marker score thresholds, accounting for gene expression variability across cell types.

To ensure the robustness of these marker scores, *genular* employs a bootstrapping approach, generating 250 resamples of the dataset (Fig. 2b). For each resample, the 10th percentile of gene expression is calculated, providing a distribution of marker score estimates. This process generates a distribution of percentile estimates for each gene across all resamples, allowing *genular* to establish confidence intervals, typically at the 95% level, around the estimated marker scores. These confidence intervals provide insights into the statistical significance and stability of the scores.

Dynamic thresholding is then utilized to set adaptive thresholds based on the bootstrapped distributions, allowing the platform to account for the inherent variability and distributional characteristics of each gene’s expression data (Fig. 2c). A gene is designated as a marker for a cell type if its expression deviation exceeds the dynamically determined threshold within that cell type, ensuring that only genes with statistically significant and biologically relevant expression changes are selected as markers. To facilitate comparison across different genes and cell types, marker scores are normalized by scaling the deviation values to a standardized range, typically between 0 and 1. This normalization accounts for factors such as gene-specific expression variability and cell type abundance, mitigating potential biases and ensuring equitable representation of marker scores.

The normalized marker scores are further integrated with gene ontology (GO) annotations, functional pathways, and protein interaction networks, enriching the marker scores with biological context. This integration allows for the identification of functionally related gene sets and pathways that are characteristic of specific cell types. *genular* performs enrichment analyses to determine whether specific GO terms or pathways are overrepresented among the high-scoring marker genes, providing deeper biological insights into the functional roles of the identified markers. Each gene receives a final marker score that encapsulates its expression deviation, statistical significance from bootstrapping, and functional relevance from ontology mapping. These comprehensive scores are used to rank genes in terms of their suitability as cell type-specific markers, categorizing them into high-confidence markers, potential markers, and non-markers based on dynamic thresholds and confidence intervals.

By exploring *genular*, we visualize the gene expression profile at the cell type level (Fig. 3a) and pathway-level distribution of gene expression at the cell level (Fig. 3b). *genular* captures cellular heterogeneity across various biological pathways, including metabolism, signal transduction, and immune system functions.

**Fig 3.**
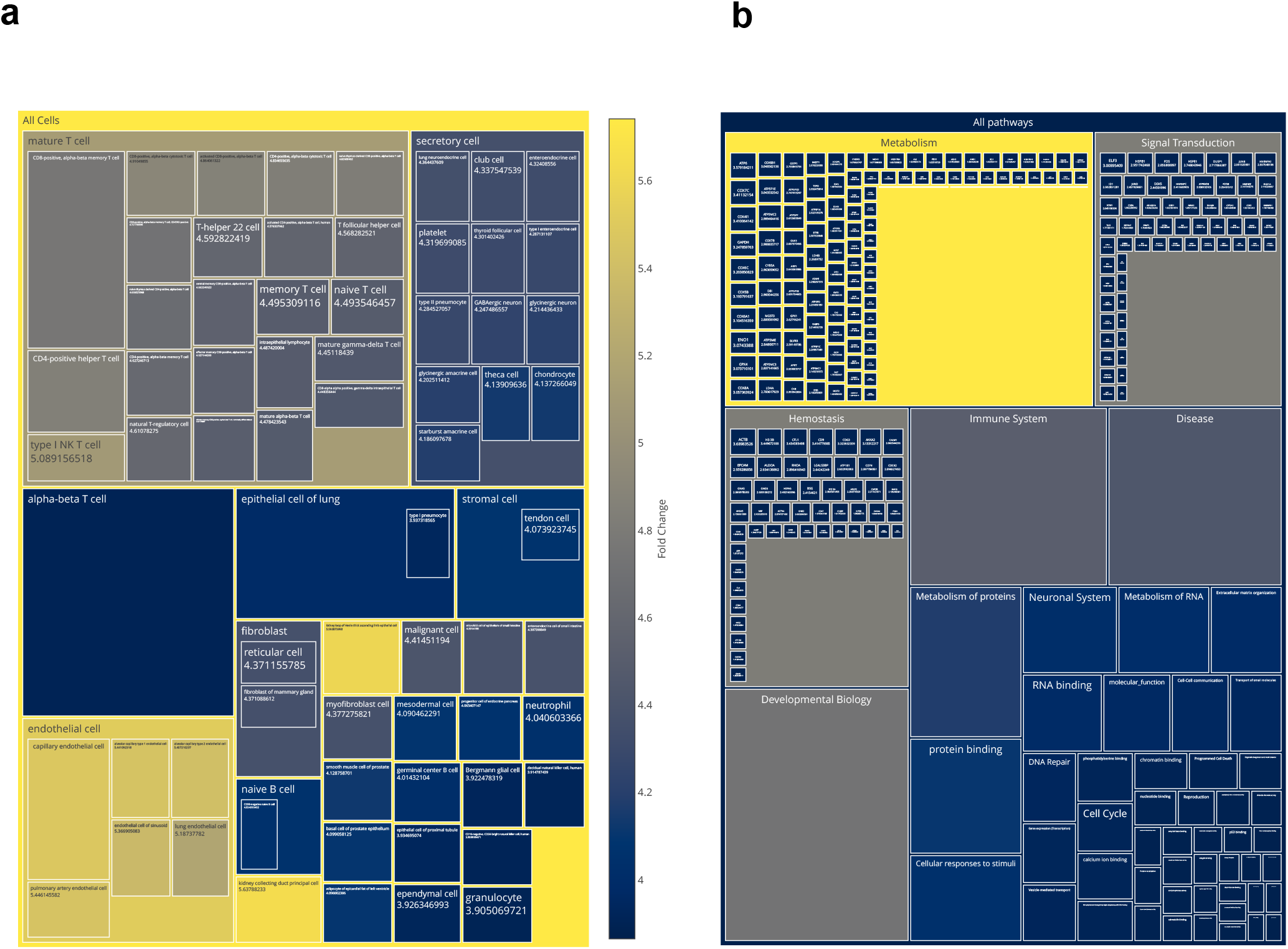
Visualization of gene expression profiles. (a) Expression of actin beta (ACTB) across various cell types, highlighting its distribution and variation. (b) Pathway-level distribution of gene expression across epithelial cells, illustrating the involvement of key biological pathways in cellular functions.

### genular Identifies Gene Signatures for Insights into Cellular Function, Differentiation, and Tissue-Specific Roles

#### Cell-type-Specific Gene Signatures Define Immune Cell Subtypes

We conducted an in-depth analysis of gene expression profiles obtained using the *genular* platform for macrophages (CL0000235), conventional dendritic cells (cDCs; CL0000990), cytotoxic T lymphocytes (CTLs; CL0000794), and helper T cells (CL0000492) due to their distinct roles in innate and adaptive immunity. By leveraging marker score calculations, we identified genes significantly enriched in each cell type, facilitating a comparative examination of their genetic landscapes (Fig. 4). Hierarchical clustering of the top 100 marker genes revealed genetic proximities aligning with immune functions: CTLs clustered with helper T cells, and macrophages with cDCs (Fig. 4a). Ubiquitously expressed genes such as ACTB, B2M, CALM1, and SLC25A5 were identified across all cell types, indicating shared roles in cytoskeletal organization, antigen processing, calcium signaling, and metabolic processes. cDCs exhibited the highest *genular* marker scores for C4B (2.21), C4A (2.20), ALPG (2.20), PHOX2A (2.20), ART4 (2.20), and AVPR2 (2.20) (Fig. 4b). C4A and C4B encode complement system components critical for immune complex clearance and opsonization, facilitating antigen presentation to T cells[20]. Reactome pathway enrichment analysis highlighted significant involvement in clathrin-mediated endocytosis and activation of C3 and C5, underscoring cDCs’ role in antigen uptake and initiation of adaptive immunity[51]. Macrophages showed high marker scores for C1QA (1.47), RHOB (1.46), C1QC (1.39), and C5AR1 (1.20) (Fig. 4c). C1QA, C1QB, and C1QC form the C1 complex, initiating the classical complement pathway essential for pathogen recognition and clearance via phagocytosis[35]. RHOB regulates actin cytoskeleton organization, crucial for phagosome formation and macrophage mobility[59]. Additionally, analysis revealed macrophage-specific expression of RUNX1, a member of the RUNX family of transcription factors involved in macrophage differentiation from monocytes[44], which has been further confirmed by the pathway enrichment analysis revealing pathways like Regulation of Complement Cascade and RUNX1-mediated transcription regulation. This comparison demonstrates that the distinct *genular* marker scores correspond to functional differences between cDCs and macrophages. While cDCs specialize in antigen uptake and presentation to T cells via clathrin-mediated endocytosis, macrophages focus on phagocytosis and pathogen clearance, supported by high expression of C1QA/B/C and involvement in phagosome formation pathways.

**Fig 4.**
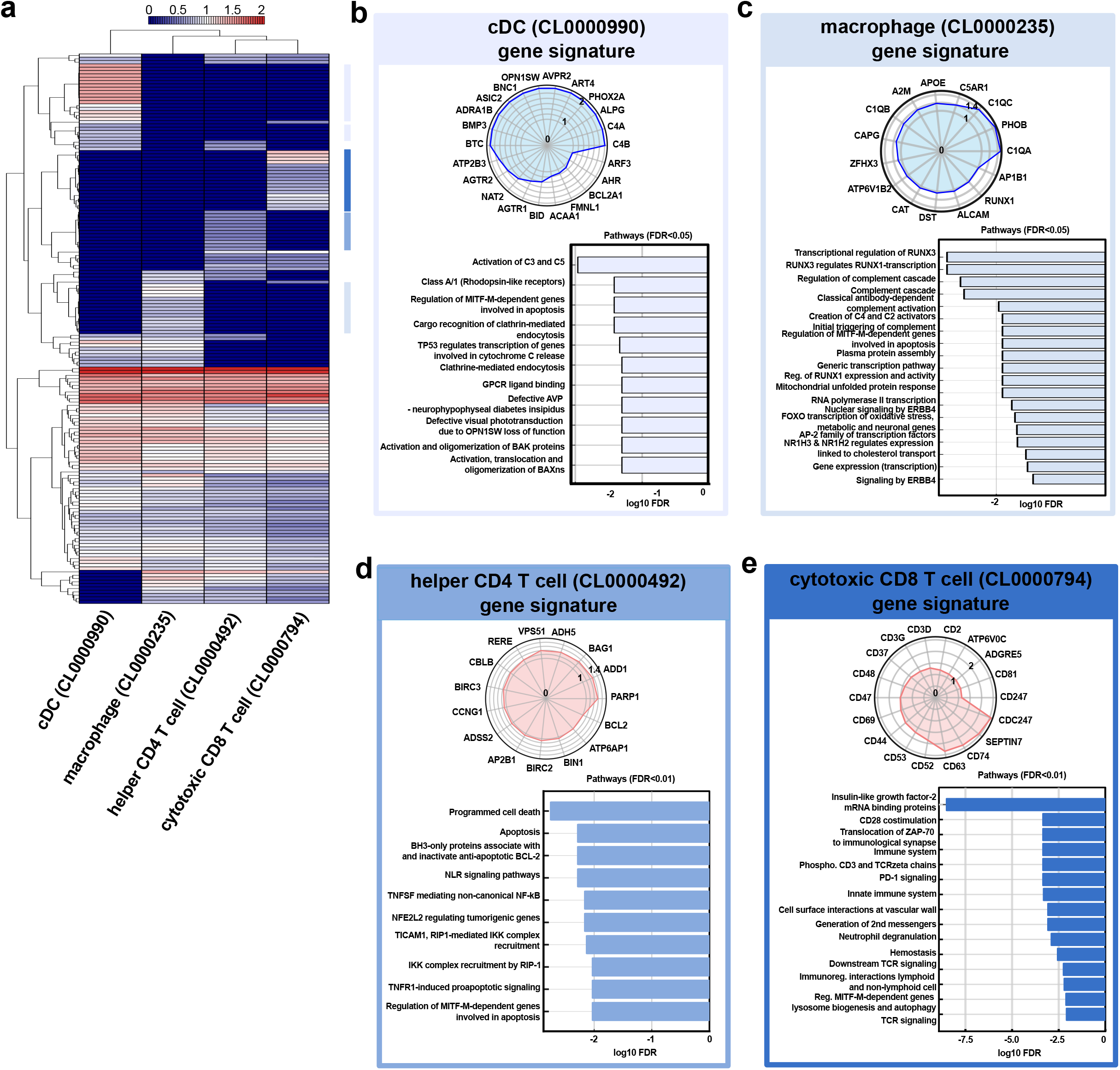
Identification of gene signatures for immune cell subtypes. (a) Hierarchical clustering of the top 100 marker genes across conventional dendritic cells (cDCs), macrophages, helper T cells, and cytotoxic T cells, highlighting shared and unique gene signatures. (b) Radial plot and bar plot showing cDC-specific gene signatures and pathway enrichment (FDR *<* 0.05). (c) Macrophage-specific gene signatures with associated enriched pathways (FDR *<* 0.05). (d) Helper T cell-specific genes and pathways enriched at FDR *<* 0.01. (e) Cytotoxic T cell-specific genes and pathways enriched at FDR *<* 0.01.

Using *genular* marker scores, we identified key genes that distinguish helper T cells from CTLs, validating the biological relevance of this approach. Helper T cells exhibited high marker scores for PARP1 (1.29), ADD1 (1.27), BAG1 (1.23), and BIRC3 (1.08) (Fig. 4d). These genes are associated with DNA repair, cytoskeletal organization, and anti-apoptotic functions that promote cell survival and proliferation—processes essential for orchestrating immune responses[5, 36, 56].

In contrast, CTLs showed elevated scores for CDC42 (2.56), SEPTIN7 (2.44), CD74 (2.37), and CD63 (2.35) (Fig. 4e). These genes are involved in actin cytoskeleton dynamics, immunological synapse formation, and antigen processing, reflecting CTLs’ specialization in direct cytotoxic activity[69, 39]. The high expression of CDC42 and SEPTIN7 supports the need for rapid cytoskeletal rearrangements during target cell engagement, while CD74 and CD63 enhance antigen recognition and presentation capabilities. This direct comparison demonstrates that the distinct marker scores calculated using the *genular* platform correspond to the functional differences between helper T cells and CTLs. Helper T cells prioritize genes that support survival and immune regulation, whereas CTLs emphasize genes facilitating cytoskeletal changes and cytotoxic functions. Reactome pathway enrichment analysis corroborated these findings: helper T cells were enriched in pathways related to programmed cell death and NLR signaling, aligning with their role in immune regulation and maintenance; CTLs were enriched in T-cell receptor signaling and PD-1 signaling pathways, consistent with their activation and effector functions in adaptive immunity.

These results confirm that *genular* marker scores effectively capture biologically meaningful gene expression differences between immune cell types. The alignment of marker scores with known functional roles underscores the validity of this metric and supports its use in identifying cell type-specific genes, enhancing our understanding of immunological processes.

#### T Cell State-Specific Gene Expression Reveals Distinct Functional States

Using *genular* marker scores, we analyzed T cells in different functional states—naive, effector, central memory (CM), and effector memory (EM)—to determine the effectiveness of our platform in distinguishing these states based on gene profiles. Hierarchical clustering of the top 100 genes revealed that effector and EM T cells clustered together, reflecting their shared roles in rapid immune responses and cytotoxic functions (Fig. 5a). In contrast, naive and CM T cells formed a separate cluster, consistent with their roles in immune surveillance and readiness for activation[26, 16, 53, 34].

**Fig 5.**
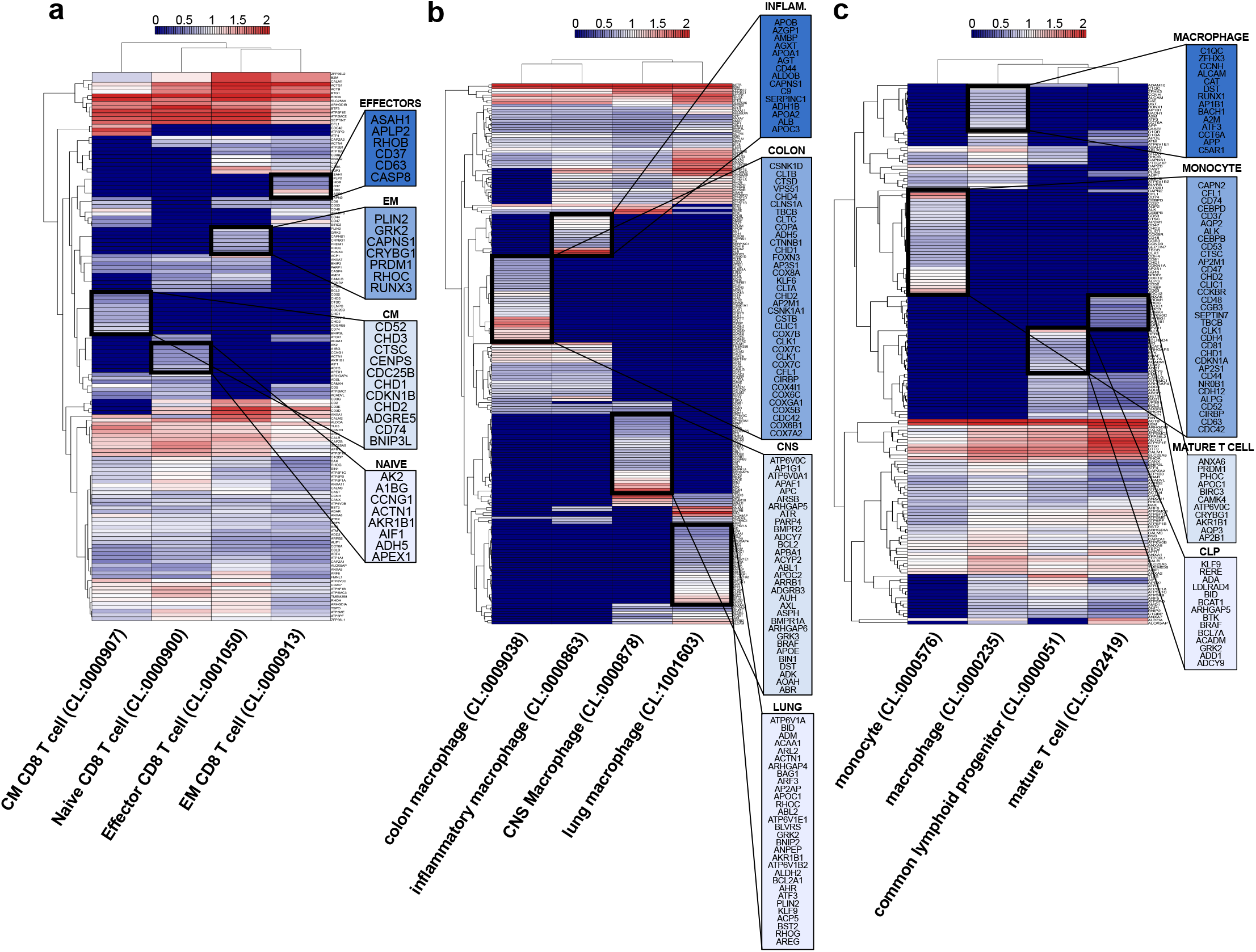
Gene expression dynamics across T cell states, macrophage tissue adaptations, and differentiation pathways. (a) Differential gene expression profiles of T cells in various states, including naive, effector, central memory (CM), and effector memory (EM) cells. (b) Tissue-specific gene expression adaptations of macrophages from the lung, colon, central nervous system (CNS), and inflammatory states, highlighting functional plasticity. (c) Gene expression changes during the differentiation pathways of monocytes to macrophages and lymphoid progenitors to T cells.

Naive T cells exhibited high *genular* marker scores for genes such as ACAA1, AK2, ACTN1, and CCNG1, which are associated with metabolic quiescence and cytoskeletal organization. ACAA1 is involved in fatty acid *β*-oxidation, contributing to energy production in resting cells[32]. AK2 plays a role in nucleotide metabolism essential for cell maintenance[42]. CCNG1 regulates the cell cycle, maintaining naive T cells in a quiescent state[1].

CM T cells showed elevated *genular* marker scores for BCL2, CD52, CHD1, and CDKN1B. BCL2 is an anti-apoptotic gene promoting cell survival, which is vital for long-lived memory cells[6]. CD52 is involved in T cell activation and signaling[3]. CDKN1B encodes a cyclin-dependent kinase inhibitor that regulates cell cycle progression, contributing to the maintenance of memory cells[67].

Effector T cells displayed increased marker scores for PLIN2, BIRC3, PRDM1, and RHOC, genes involved in apoptosis inhibition, lipid metabolism, terminal differentiation, and cytoskeletal dynamics essential for effector functions[52]. PRDM1 encodes BLIMP-1, a transcription factor critical for terminal differentiation of effector cells[24]. EM T cells showed high scores for AQP3, ASAH1, CD37, and CD63, which facilitate cell migration, influence cell signaling and apoptosis, and are critical for immune synapse formation and vesicular trafficking necessary for rapid effector responses[19, 18, 46].

These findings demonstrate that *genular* marker scores effectively distinguish between T cell functional states, capturing biologically meaningful gene expression differences that correspond to their known immunological roles.

#### Tissue-Specific Adaptations of Macrophages Highlight Functional Plasticity

We further applied *genular* marker scores to macrophages from different tissues—the colon, central nervous system (CNS), lung, and inflammatory states—to assess whether the marker scores calculated using our platform can differentiate macrophage tissue specificity based on gene expression profiles. Clustering analysis showed that lung and CNS macrophages clustered together, reflecting shared roles in tissue surveillance and immune regulation in specialized environments, while inflammatory and colon macrophages formed a separate cluster, indicating their involvement in active immune responses (Fig. 5b).

Colon macrophages exhibited high marker scores for genes such as CLTB, CTSD, COX8A, and COX6C, associated with increased mitochondrial activity and protein degradation, reflecting the energy demands in a high-antigen environment[58]. Inflammatory macrophages showed elevated scores for APOB, CD44, SERPINC1, and AGT, involved in lipid metabolism, cell adhesion, coagulation, and inflammatory responses[41, 28]. CNS macrophages (microglia) displayed high marker scores for APOE, BMPR2, ATR, and BIN1, genes associated with neuroimmune regulation and homeostasis[73, 33, 10]. Lung macrophages exhibited elevated scores for ANXA1, ALOX5AP, PLIN2, and BCL2A1, highlighting roles in inflammation resolution and pathogen response[45, 30]. These results indicate that *genular* marker scores effectively distinguish macrophages based on tissue-specific gene expression profiles, highlighting their functional plasticity and adaptation to the tissue microenvironment.

#### Gene Expression Dynamics During Differentiation

We explored the monocyte-to-macrophage transition and lymphoid progenitor-to-T cell differentiation, revealing stage-specific gene expression changes. Hierarchical clustering of the top marker genes showed a close genetic similarity between lymphoid progenitors (CL0000051) and T cells (CL0002419), indicating a direct differentiation pathway. In contrast, monocytes (CL0000576) clustered closely with macrophages (CL0000235), reaffirming their established lineage relationship and distinct differentiation trajectory (Fig. 5c). Focusing on the monocyte-enriched gene set, several key regulatory and functional genes were identified. Transcription factors such as CEBPB and CEBPD play prominent roles in maintaining the monocyte state by regulating genes involved in cell proliferation, differentiation, and immune responses[22]. Signaling molecules including CD74 and CD47 were highly expressed, highlighting their involvement in antigen presentation and cell-cell interactions essential for monocyte function[43].

In contrast, the macrophage-enriched gene set revealed a distinct set of regulatory and functional genes critical for macrophage specialization and function. The transcription factor RUNX1 emerged as a key regulator, orchestrating the expression of genes involved in macrophage activation, differentiation, and response to inflammatory stimuli[44]. Signaling molecules like C5AR1 and AIF1 were prominently featured, emphasizing their roles in inflammatory signaling and phagocytosis. Overall, in the monocyte-to-macrophage pathway, genes involved in phagocytosis and homeostasis, such as RUNX1 and C5AR1, were progressively upregulated with the downregulation of monocyte-associated genes, such as CEBPB and CD74. This shift signifies a reprogramming event that transitions cellular functions from maintaining a proliferative and migratory monocyte state to adopting specialized roles in immune defense, tissue remodeling, and homeostasis.

Comparative analysis of lymphoid progenitors and T cells demonstrated a clear genetic switch, characterized by the downregulation of progenitor-associated genes such as KLF9 and RERE (crucial for T cell lymphopoiesis[72]) and the upregulation of T cell-specific genes, including PRDM1 and CAMK4. The transcription factor PRDM1 emerged as a pivotal master regulator, orchestrating the repression of progenitor-specific genes while activating T cell-specific programs[71]. This transcriptional reprogramming is essential for driving the maturation process from progenitor to fully differentiated T cells. Metabolic enzymes, including APOE, PLIN2, and AKR1B1, were significantly upregulated in T cells, reflecting a metabolic shift towards pathways that support the high energy and biosynthetic demands of active T cells. This metabolic reprogramming facilitates rapid proliferation and the execution of effector functions necessary for effective immune responses. This shift signifies a reprogramming event that reorients cellular functions from maintaining a proliferative progenitor state to adopting specialized immune functions.

Overall, *genular* provides insights into the regulatory networks, signaling pathways, and metabolic adaptations that drive essential immune functions by successfully delineating the genetic landscapes of T cell maturation and the monocyte-to-macrophage transition. The ability to uncover and analyze key transcription factors, signaling molecules, and metabolic enzymes highlights *genular*’s capacity to integrate complex single-cell transcriptomic data with comprehensive biological databases. In summary, *genular* is a powerful tool for advancing our understanding of cellular functions, differentiation mechanisms, and tissue-specific roles, offering valuable implications for therapeutic strategies and biomedical research.

## Discussion

*genular* represents a significant advancement in the field of single-cell transcriptomics by addressing critical limitations of existing platforms through its innovative integration of large-scale scRNA-seq data with comprehensive genomic and proteomic information. A central feature of *genular* is its ability to calculate a cell marker score for each gene, enabling the precise quantification of gene expression across cells. This facilitates the derivation of unique profiles specific to cell types, states, and lineages, thereby enhancing the resolution and accuracy of cellular identity definitions. Current state-of-the-art tools often struggle with the sheer volume and complexity of scRNA-seq data, relying heavily on computationally intensive clustering methods like t-SNE or UMAP that can obscure subtle gene expression differences. In contrast, *genular* ‘s novel statistical framework bypasses these limitations by directly quantifying gene expression deviations relative to average levels across cell types. This approach not only improves computational efficiency but also enhances the biological interpretability of the results, allowing for the identification of both shared and cell-type-specific gene signatures with greater precision. Using *genular*, we successfully determined T cell memory states, mapped differentiation profiles by tracking gene expression changes as monocytes mature into macrophages and lymphoid progenitor cells develop into T cells, and captured tissue-specific reprogramming of macrophages. These applications demonstrate *genular* ‘s capability to reveal distinct gene expression profiles that underpin specialized functions in different tissues. Such detailed insights are critical for understanding immune cell functions and their roles in various physiological and pathological contexts, thereby closing gaps in our ability to characterize cellular heterogeneity and dynamic gene regulation. Furthermore, *genular* ‘s integrated database architecture, built on MongoDB, harmonizes data from 16 established biological repositories, including NCBI Gene, Human Protein Atlas, STRING, and UniProt. This comprehensive integration ensures that gene functions, pathways, and protein interactions are seamlessly connected to gene expression profiles, providing a holistic view of cellular identities and functions. Unlike existing platforms that may operate in silos or struggle with data interoperability, *genular* ‘s unified framework facilitates cross-study comparisons and scalable analyses, making it a versatile tool for large-scale single-cell investigations. Looking forward, we envision *genular* evolving into a community-driven platform that continuously incorporates new omics data types, such as proteomics and epigenomics, to provide an even more comprehensive view of cellular states and regulatory mechanisms. The open-source nature of *genular*, combined with its flexible architecture, encourages contributions from the scientific community, fostering collaborative advancements and ensuring the platform remains at the forefront of single-cell analysis technologies. Additionally, integrating advanced machine learning techniques could further enhance *genular*’s predictive capabilities, enabling dynamic modeling of cellular interactions within complex microenvironments and facilitating the discovery of novel regulatory networks. In conclusion, *genular* bridges critical gaps in single-cell transcriptomic analysis by offering a scalable, integrated, and efficient platform that enhances the accuracy and depth of cellular identity and function characterization. Its innovative marker score calculations and comprehensive data integration set a new standard in the field, providing researchers with the tools necessary to uncover nuanced biological insights and accelerate discoveries in immunology, gene regulation, cellular differentiation, and disease research. As single-cell datasets grow in volume and diversity, *genular* is well-positioned to support cutting-edge research, driving our understanding of complex biological systems forward and contributing to the development of targeted therapies and personalized medicine.

## Conclusion

*genular* serves as a robust and comprehensive open-source platform for the integration, exploration, and analysis of genomic and proteomic data. Its scalable architecture, coupled with flexible access options, positions it as a vital tool for researchers in the biological sciences, enabling more profound insights into complex biological systems.

We actively welcome contributions from the scientific community and encourage researchers to engage in the continued development and enhancement of *genular*, ensuring it remains a valuable resource for advancing biomedical research. The platform’s source code and documentation are available on GitHub at https://github.com/atomiclaboratory/genular-database.

## Data Availability Statement

All data used in this manuscript, including gene annotations, protein sequences, and gene expression data, are publicly available from established resources such as NCBI Gene [40], Human Protein Atlas [64], STRING [61], Reactome [37], and CELLxGENE [13].

The primary datasets were retrieved using automated scripts, and processed data, including effect sizes and marker scores, are available through the Genular platform [63]. The Genular platform provides open access to the integrated database via a RESTful API, an R package [27], and a downloadable MongoDB dump [23], which can be accessed at https://genular.atomic-lab.org. Specific instructions for data access and API usage are provided in the platform’s documentation.

## Code Availability

The code used to process the datasets and generate the results for this manuscript is publicly available. The Genular database source code can be accessed on GitHub at https://github.com/atomiclaboratory/genular-database, which includes both the API and R package [27] repositories. Additionally, the scripts used for data retrieval and analysis are available upon request or via the linked repositories.

## Supplementary Methods

### R Package Usage Examples

The *genular* R package provides powerful tools to integrate single-cell datasets into R-based workflows. Below, we present several practical examples of how to leverage this package to retrieve and analyze data from the *genular* database.

Example 1: Determining Cell IDs and Marker Score Thresholds

The following R code example demonstrates how to query specific cell types from the *genular* database and retrieve their associated marker scores, along with search scores.

**Listing 1**. R Code Example: Retrieving Cell IDs and Marker Score Thresholds

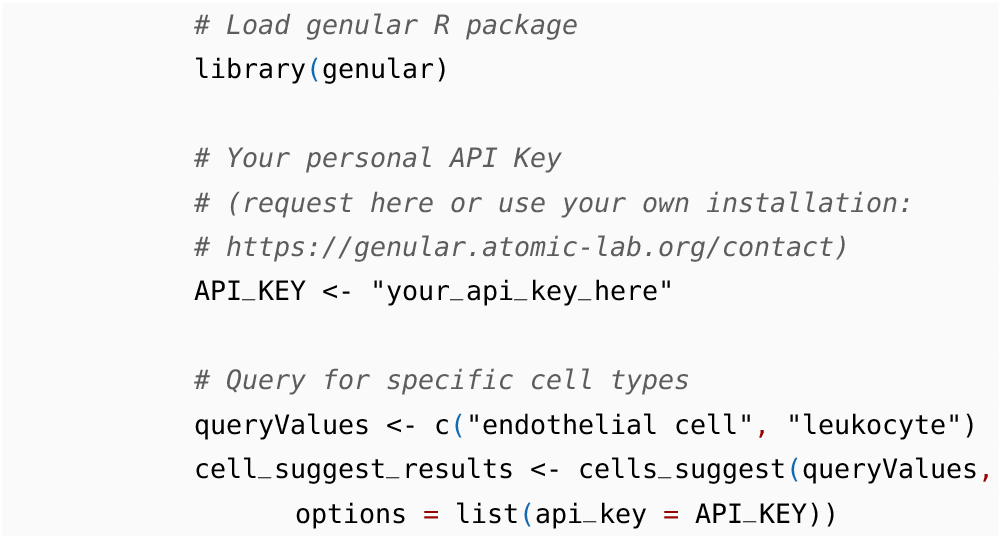

The JSON output contains cell IDs, names, search scores, and marker scores for the queried cell types:

**Listing 2**. Example Output: JSON Response for Cell Search

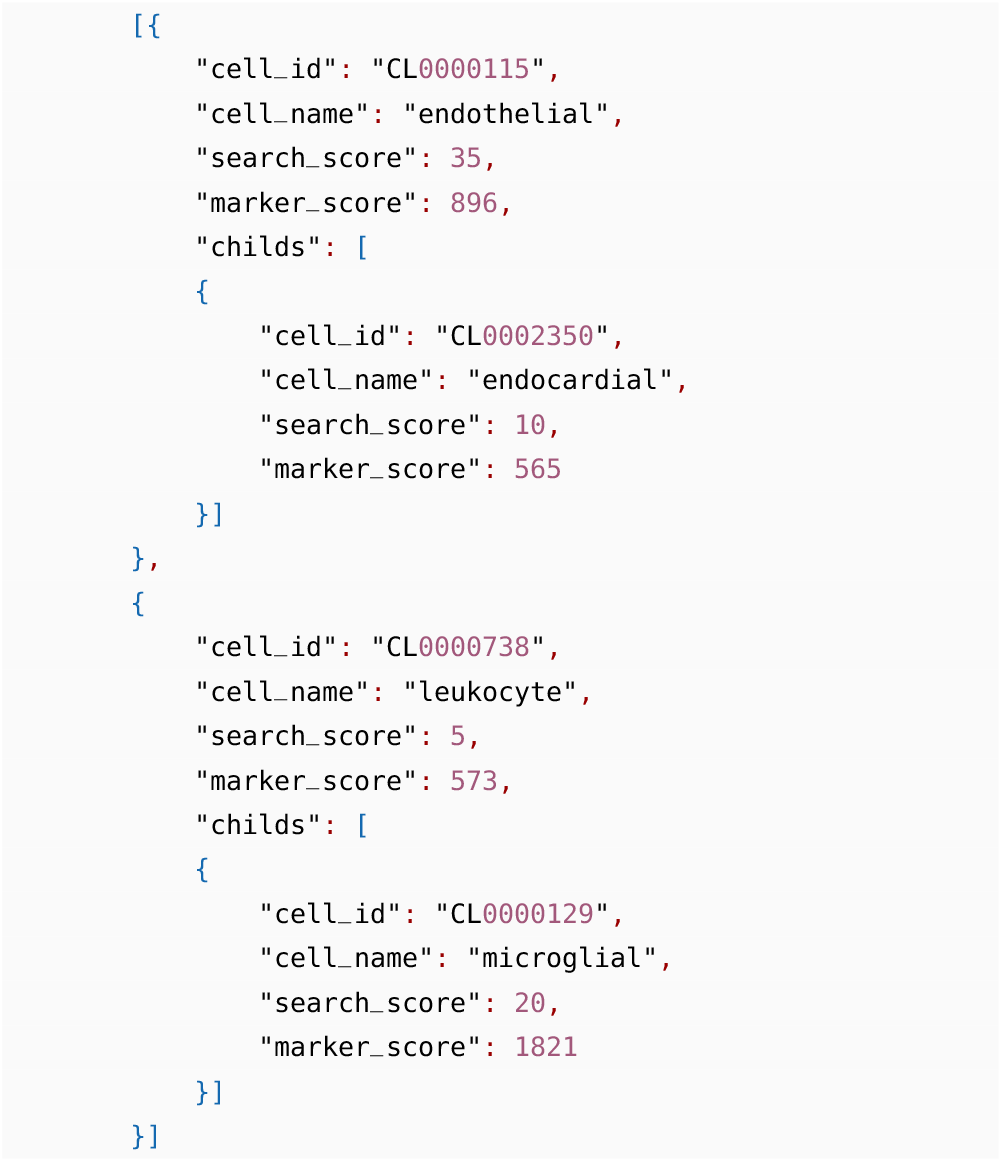

This output represents the results from querying for cell IDs and calculating marker score thresholds.

Example 2: Retrieving Single-Cell Gene Expression Profiles Across Specific Cell Types

The following R code allows users to query the *genular* database for gene expression profiles across specific cell types, including marker scores and effect sizes.

**Listing 3**. R Code Example: Querying Gene Expression Profiles Across Specific Cell Types

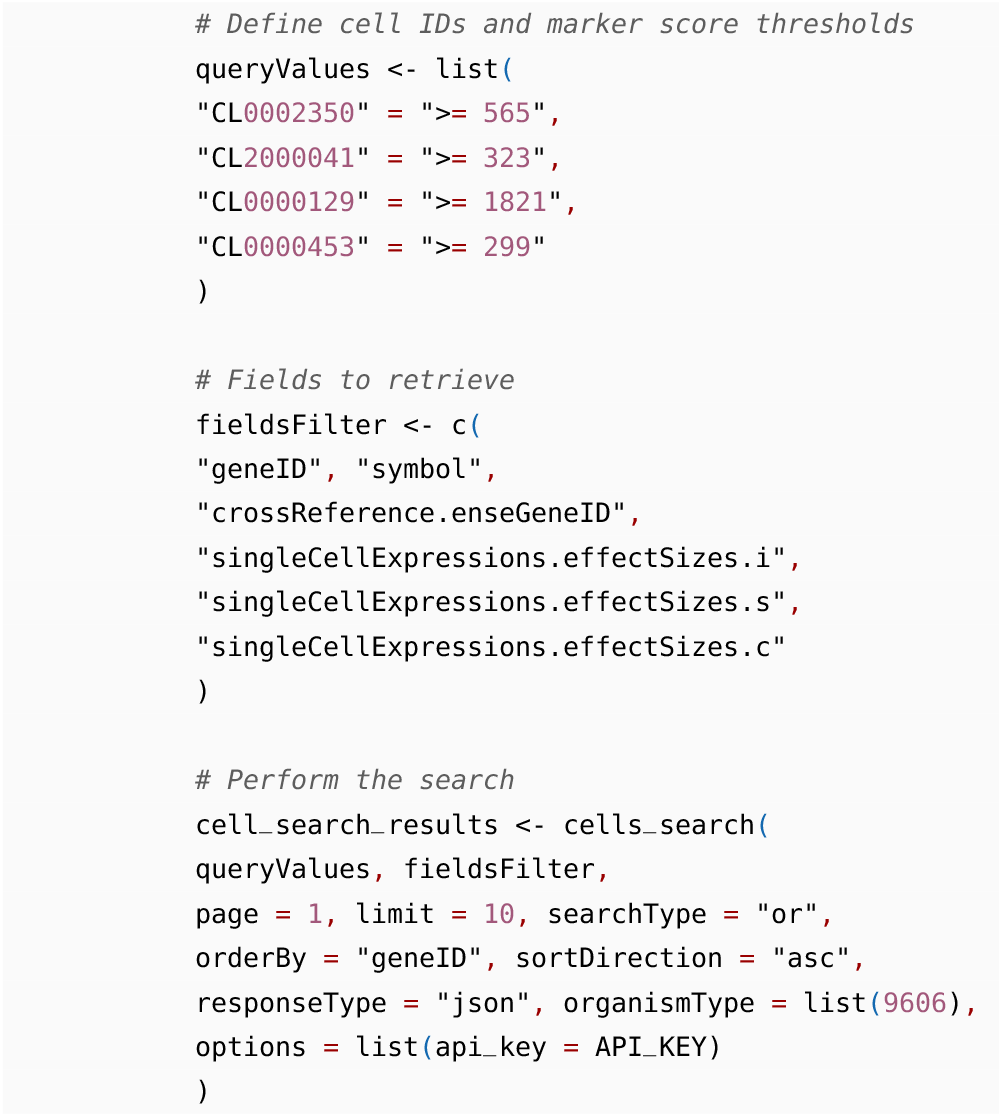

This example provides gene expression profiles, marker scores, and score thresholds for the queried cell types:

**Listing 4**. Example Output: JSON Response for Gene Expression Profiles and Marker Scores

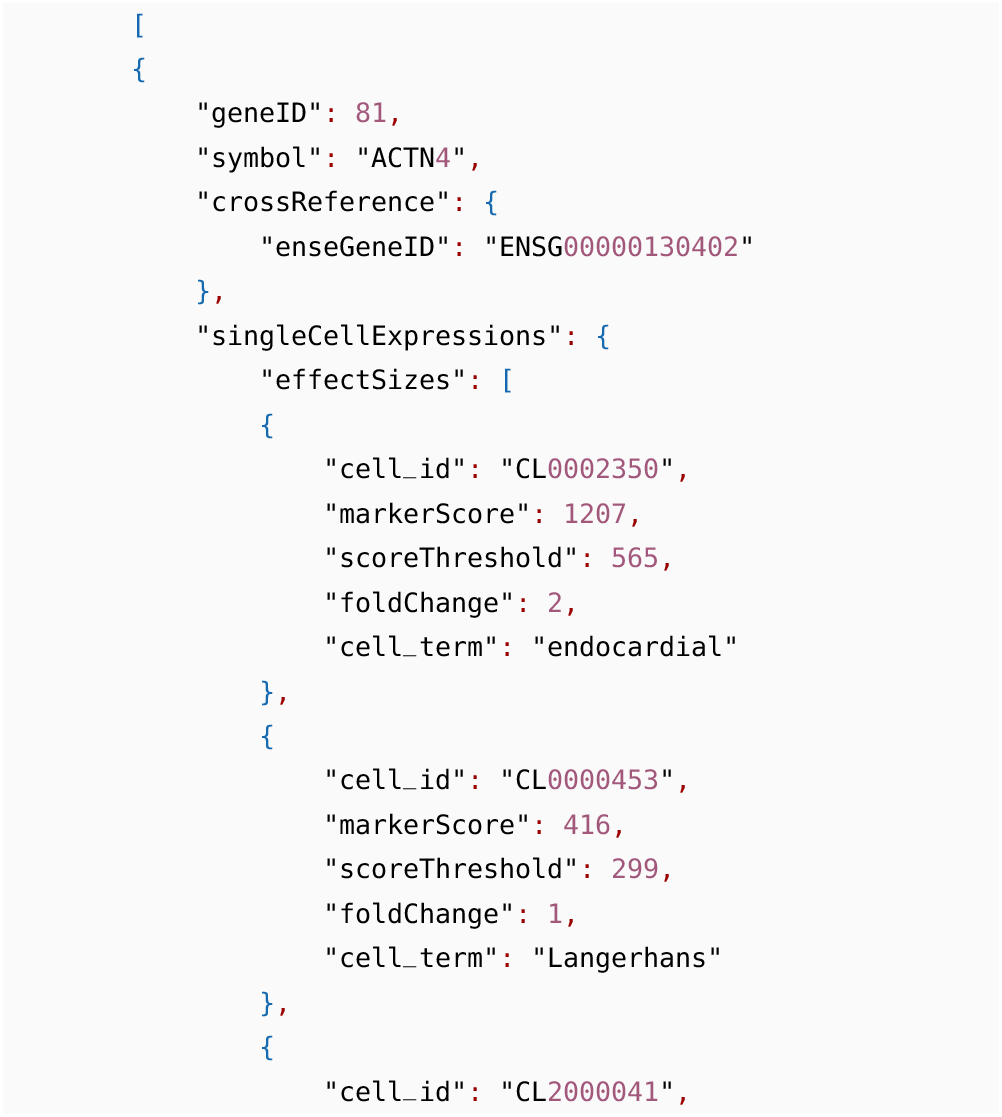
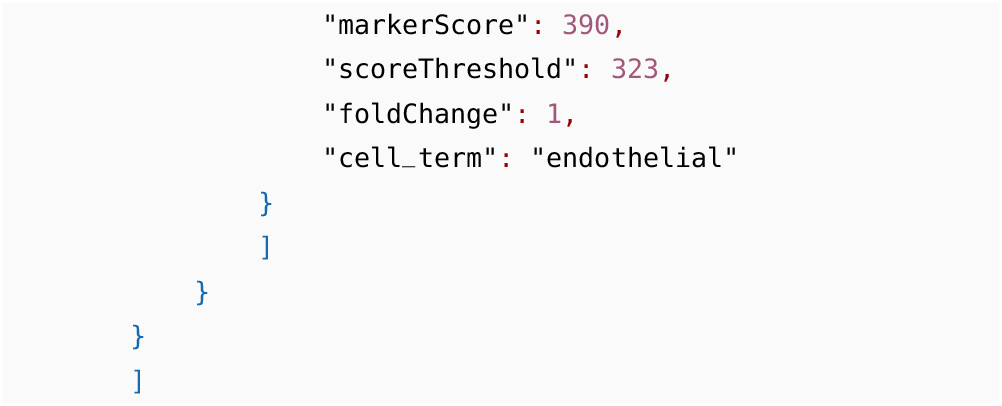

Example 3: Searching for Gene Information

The following R code enables users to search for specific genes and retrieve relevant gene-specific data, including gene symbols, Ensembl gene IDs, and effect sizes across cell types.

**Listing 5**. R Code Example: Retrieving Gene-Specific Information

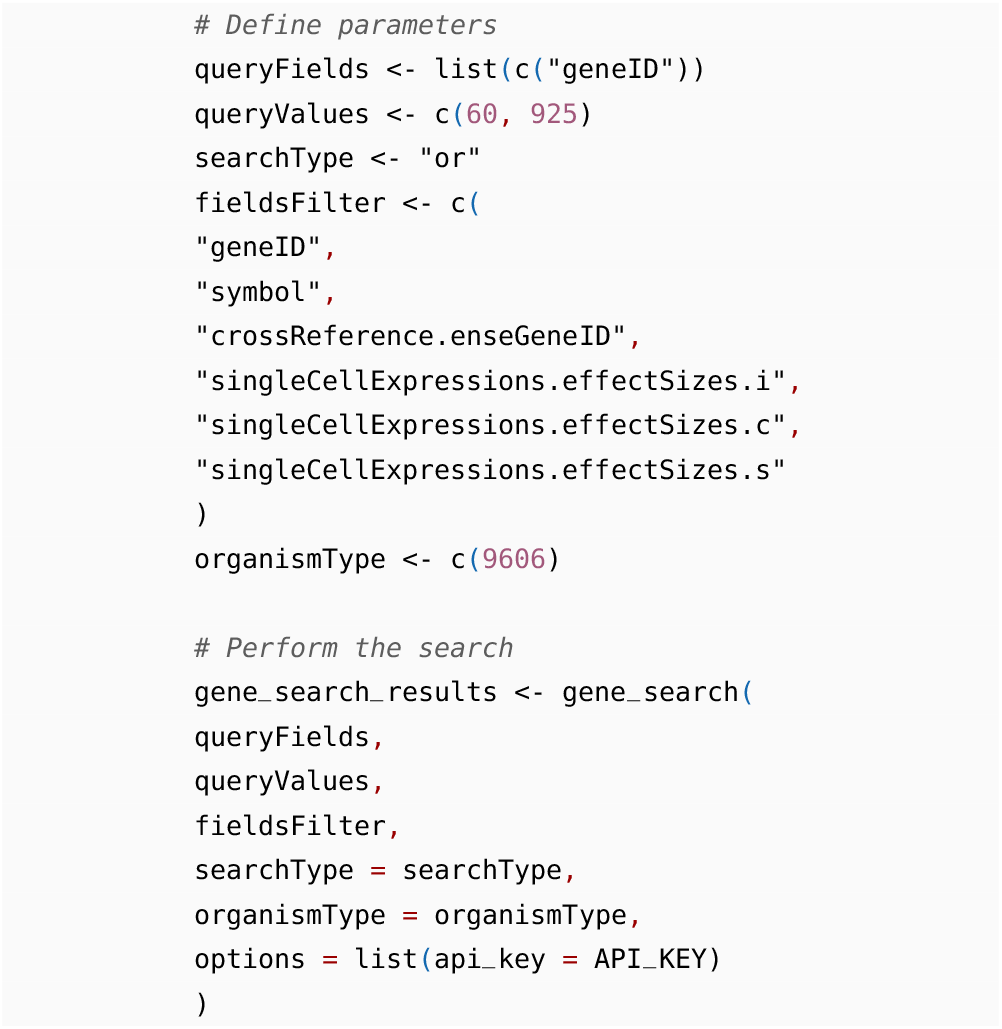

The JSON response contains detailed gene information and the effect sizes for each gene across cell types:

### Cell Lineage Statistics for Top Unique Cells

This section provides a summary of the top cell lineages where the unique cell count exceeds 500,000, highlighting the diversity and abundance of these cell types within the Genular database. The table below lists the cell lineages, including their corresponding unique cell counts, demonstrating the scale of data coverage for key immune, endothelial, neuronal, and other cell types.

|  | Cell ID | Cell Name | Unique Cell Count |
| --- | --- | --- | --- |
| 1 | CL0000115 | endothelial cell | 1658215 |
| 2 | CL0000057 | fibroblast | 1631465 |
| 3 | CL0000235 | macrophage | 1416841 |
| 4 | CL0000236 | B cell | 1358536 |
| 5 | CL0000084 | T cell | 1152857 |
| 6 | CL0000097 | mast cell | 1092282 |
| 7 | CL0000669 | pericyte | 1064328 |
| 8 | CL0000786 | plasma cell | 1064137 |
| 9 | CL0000623 | natural killer cell | 849449 |
| 10 | CL0000576 | monocyte | 805587 |
| 11 | CL0000451 | dendritic cell | 802700 |
| 12 | CL0000540 | neuron | 796395 |
| 13 | CL0000625 | CD8-positive, alpha-beta T cell | 777413 |
| 14 | CL0000763 | myeloid cell | 745394 |
| 15 | CL0000624 | CD4-positive, alpha-beta T cell | 720814 |
| 16 | CL0000160 | goblet cell | 717580 |
| 17 | CL0000128 | oligodendrocyte | 639251 |
| 18 | CL0000066 | epithelial cell | 636233 |
| 19 | CL0002453 | oligodendrocyte precursor cell | 601739 |
| 20 | CL0000646 | basal cell | 598707 |
| 21 | CL0000127 | astrocyte | 587930 |
| 22 | CL0002138 | endothelial cell of lymphatic vessel | 569786 |
| 23 | CL0000814 | mature NK T cell | 561759 |
| 24 | CL0000815 | regulatory T cell | 535409 |
| 25 | CL0000860 | classical monocyte | 524346 |
| 26 | CL0000875 | non-classical monocyte | 519532 |
| 27 | CL0000784 | plasmacytoid dendritic cell | 511624 |
| 28 | CL0000192 | smooth muscle cell | 509476 |
| 29 | CL0000738 | leukocyte | 502744 |

## Contributions

I.T. led the software development for the *genular* database, overseeing its architecture, design, and implementation. A.T., provided critical intellectual contributions in the project’s conceptual framework, along with securing funding and offering domain-specific expertise in the field of biomedical research. S.H., made substantial contributions to the project, including developing key use case examples, generating visualizations, and offering insights into the application of *genular* in various research contexts. The success of this project reflects the collective efforts and collaboration of the entire team.

